# EES-Transformer: A Dual-Path Transformer for Tissue Classification and Gene Representation Learning from Extreme Expression Sets

**DOI:** 10.64898/2026.02.22.707241

**Authors:** Joo-Seok Park, Yejin Lee, Yang Jae Kang

## Abstract

Accurate tissue classification from gene expression data is fundamental to transcriptomic analysis. Here we introduce EES-Transformer V2, a dual-path transformer architecture that learns from Extreme Expression Sets (EES)—sequences of genes at expression extremes (above 95th or below 5th percentile). The architecture separates tissue classification from masked language modeling through independent branches: the classification branch operates without tissue label information, while the generative branch receives tissue conditioning. This design enables fair evaluation of classification performance while learning tissue-specific gene relationships. Applied to 12,212 Arabidopsis thaliana RNA-seq samples spanning 47 tissue types, EES-Transformer achieves 91-92% classification accuracy (varying across evaluation runs due to stochastic input masking)—substantially above the 2.1% random baseline. Attention-based analysis identifies 3,524 gene-tissue high-attention associations whose importance patterns reflect known biology. Critically, while individual high-attention genes appear broadly across tissues, gene pairs from attention-derived regulatory networks show higher tissue specificity: pollen gene pairs show a 2.7-fold enrichment over single-gene rates, and root and leaf pairs each show 1.5-fold enrichment. This finding reveals that tissue identity is encoded in combinatorial gene expression patterns rather than individual genes. Attention-derived gene regulatory networks exhibit scale-free topology and biologically coherent hub gene programs, with pollen networks consisting entirely of DOWN-DOWN interactions among silenced vegetative genes. EES-Transformer provides accurate tissue classification, interpretable gene importance scores, and attention-derived regulatory networks for biological discovery.

## Introduction

Gene expression profiling through RNA sequencing has become indispensable for understanding tissue-specific biology, developmental processes, and disease states [1]. A fundamental challenge in transcriptomics is accurately identifying the tissue of origin from expression profiles—essential for quality control, sample annotation, and biological interpretation [2]. While deep learning approaches have transformed many areas of computational biology, applying these methods to tissue classification presents unique challenges related to data representation and fair model evaluation.

Transformer architectures have demonstrated remarkable success across domains, from natural language processing to protein structure prediction [3]. Recent work has adapted transformers for gene expression analysis, treating genes as tokens and expression profiles as sequences [4,5]. These approaches typically employ masked language modeling (MLM) objectives borrowed from natural language processing, where the model learns to predict masked tokens from context.

Several approaches have been developed for expression-based tissue classification. Traditional machine learning methods, including k-nearest neighbors, random forests, and support vector machines, can achieve high accuracy when trained on sufficient data. Recent work by Palande et al. demonstrated that k-nearest neighbors classifiers achieve near-perfect precision and recall on Arabidopsis tissue classification, suggesting that expression signatures alone contain sufficient information for tissue identification [2]. However, these approaches focus solely on classification without providing interpretable gene importance scores or learned representations suitable for transfer learning. Transformer-based methods such as Geneformer [4] and scBERT [5] have shown remarkable success in cell type annotation.

We introduce three key innovations. First, we propose the Extreme Expression Set (EES) representation, which captures tissue identity through genes at expression extremes rather than continuous expression values. This discrete representation is both biologically motivated—extreme expression often reflects tissue-specific transcriptional programs—and computationally efficient, reducing each sample to a variable-length sequence of gene tokens. Second, we develop a dual-path architecture that physically separates classification and generative objectives into independent branches with distinct information flows. The tissue classification branch never observes tissue labels, while the MLM branch receives tissue conditioning to learn context-dependent gene relationships. Third, we demonstrate that attention patterns from the classification branch identify tissue-differential gene importance and, through pairwise analysis, reveal that tissue identity is encoded in combinatorial gene expression patterns.

We validate EES-Transformer on a comprehensive *Arabidopsis thaliana* dataset comprising 12,212 samples across 47 tissue types. The model achieves 91-92% classification accuracy while providing interpretable gene importance scores. Analysis of confusion patterns reveals biologically meaningful relationships between tissues, and attention-based analysis identifies high-attention genes consistent with known biology. Crucially, gene pairs from attention-derived regulatory networks are dramatically more tissue-specific than individual genes, revealing that tissue identity is encoded in combinatorial expression patterns. These results establish EES-Transformer as both a practical tool for tissue classification and a framework for biological discovery through interpretable deep learning.

## Results

### Extreme Expression Sets capture tissue-specific identity

We hypothesized that tissue identity is encoded primarily in genes at expression extremes—those exceptionally upregulated or downregulated relative to their typical expression across tissues. To test this, we developed the Extreme Expression Set (EES) representation, which converts continuous expression profiles into discrete token sequences (Fig. 1a).

**Figure 1:**
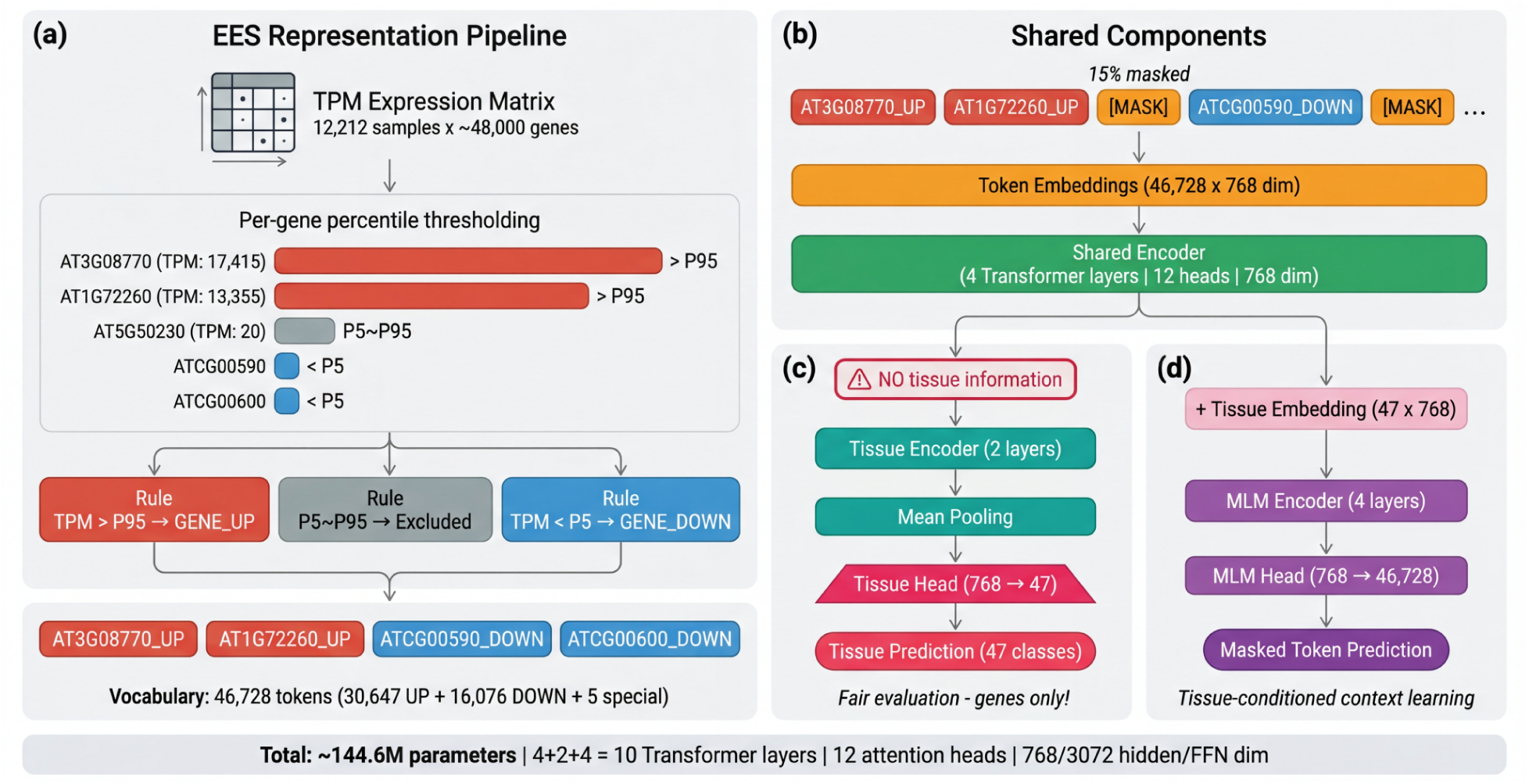
EES-Transformer V2 architecture overview. **(a)** Extreme Expression Set (EES) representation. Continuous TPM expression values are converted to discrete gene tokens by per-gene percentile thresholding. For each gene, expression percentiles are computed across all 12,212 samples. Genes with expression above the 95th percentile receive an “_UP” suffix, those below the 5th percentile receive “_DOWN”, and genes within the middle range are excluded. **(b)** Overall dual-path architecture. Input gene tokens are embedded into 768-dimensional vectors without positional encoding (permutation invariant design) and processed by a shared encoder (4 transformer layers with 12 attention heads). The shared representation then splits into two independent branches: the tissue classification branch (c) and the masked language modeling (MLM) branch (d). **(c)** Tissue classification branch. 2 additional transformer layers process the shared gene representations, followed by mean pooling across all tokens and a linear projection to 47 tissue classes. **(d)** MLM branch. Tissue embeddings (47 tissues x 768 dimensions) are added to the shared representations, conditioning the model on tissue identity. 4 transformer layers process these tissue-conditioned representations before projecting to the full vocabulary (46,728 tokens) for masked token prediction.

For each gene across all samples, we computed per-gene expression percentiles. Genes with expression above the 95th percentile in a given sample receive an “UP” designation (e.g., AT1G01010_UP), while those below the 5th percentile receive “DOWN” (e.g., AT1G01010_DOWN). This process converts each sample’s expression profile into a variable-length sequence of gene tokens, analogous to a sentence in natural language processing. After filtering genes with mean TPM below 1.0, our vocabulary comprises 46,728 tokens: 30,647 genes with UP tokens, 16,076 genes with DOWN tokens, and 5 special tokens. The asymmetry reflects the fact that not all genes reach both expression extremes across the dataset.

The EES representation offers several advantages. First, it naturally handles the high dimensionality of expression data by focusing on the most informative genes per sample. Second, the binary direction (UP/DOWN) captures whether a gene is specifically activated or repressed in a given context, rather than its absolute expression level. Third, the representation is permutation invariant—gene sets have no inherent order—which we exploit through our architecture by omitting positional encodings.

### Dual-path architecture prevents information leakage

The core innovation of EES-Transformer V2 is its dual-path design, which separates tissue classification from masked language modeling into independent branches (Fig. 1b-d). This separation ensures fair evaluation of classification performance while enabling tissue-conditioned gene representation learning.

The architecture begins with a shared encoder comprising four transformer layers with 12 attention heads and 768-dimensional hidden representations. This shared encoder learns general relationships between genes that apply regardless of the downstream task. The shared representation is then processed by two independent branches.

The tissue classification branch (Fig. 1c) receives the shared representations without any tissue information. Two additional transformer layers process these representations, followed by mean pooling across all tokens and a linear projection to 47 tissue classes. Critically, this branch never observes the tissue label—not as an input embedding, not as a conditioning signal, and not through any skip connection. Classification accuracy therefore reflects genuine learning from gene expression patterns rather than label memorization.

The MLM branch (Fig. 1d) receives both the shared representations and tissue embeddings. The tissue embedding is added to hidden states at each position, conditioning the model’s predictions on tissue identity. Four transformer layers then process these conditioned representations before a final projection to the full vocabulary. This branch learns that certain genes are expected to be highly or lowly expressed in particular tissues—knowledge that enhances biological representation learning while remaining separate from the classification objective.

The total model comprises 144.6 million parameters, with the majority residing in the embedding layer (46,728 tokens × 768 dimensions) and the MLM prediction head.

### Tissue classification achieves 91-92% accuracy across 47 tissue types

We trained EES-Transformer on 12,212 *Arabidopsis thaliana* RNA-seq samples annotated with 47 tissue types (Table S1). The model achieves 91-92% overall accuracy on held-out validation data (Fig. 2a; Figure S1), varying across evaluation runs due to stochastic 15% input masking (see Methods). This represents a ∼43-fold improvement over the 2.1% random baseline (1/47 tissues).

**Figure 2:**
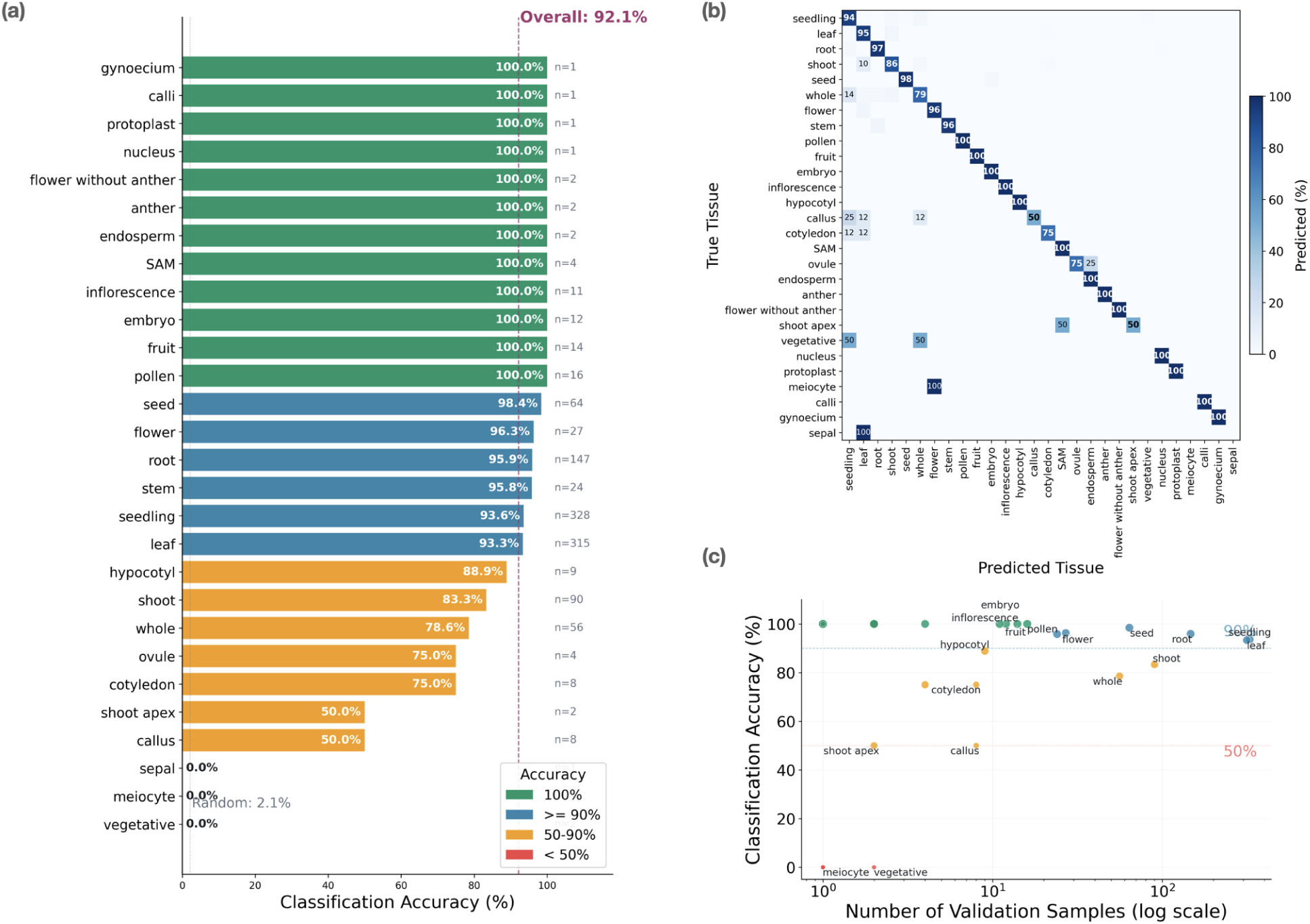
Tissue classification performance. **(a)** Per-tissue classification accuracy on the validation set (n = 1,153 samples across 28 tissue types present in validation). Horizontal bars show the percentage of correctly classified samples for each tissue type, colored by accuracy level (green: 100%, blue: >= 90%, amber: 50-90%, red: < 50%). Sample counts (n) are shown to the right of each bar. The dashed magenta line indicates overall accuracy (92.1%), and the dotted gray line shows the random baseline (2.1% = 1/47 tissues). **(b)** Row-normalized confusion matrix showing the distribution of predicted tissue labels for each true tissue type. Rows represent true tissues (sorted by sample count), and columns represent predicted tissues. Values indicate the percentage of samples from each true tissue assigned to each predicted tissue. Off-diagonal entries >= 10% are annotated, revealing biologically meaningful confusion patterns: shoot is confused with leaf (shared aerial transcriptome), whole plant with seedling (mixed tissue context), and callus with seedling (dedifferentiated expression patterns). **(c)** Relationship between validation sample size and classification accuracy. Each point represents one tissue type, with size proportional to average prediction confidence. The x-axis uses a logarithmic scale.

Per-tissue analysis reveals that accuracy varies with tissue distinctiveness and sample availability (Fig. 2a-c; Table S2). Twelve tissues achieve perfect classification, including those with unique expression signatures: pollen (100%, n=16), fruit (100%, n=14), embryo (100%, n=12), and inflorescence (100%, n=11). The model also performs well on major tissue types with abundant training data: seed (98.4%, n=64), root (95.9%, n=147), seedling (93.6%, n=328), and leaf (93.3%, n=315).

Classification is more challenging for tissues that share expression programs with related tissues or have limited training data. Shoot samples (83.3%, n=90) are frequently confused with leaf, reflecting their overlapping aerial tissue transcriptomes. Whole plant samples (78.6%, n=56) are confused with seedling, as both represent mixed tissue contexts. Cotyledon (75.0%, n=8) and callus (50.0%, n=8) show lower accuracy due to small sample sizes and, in the case of callus, dedifferentiated expression patterns that resemble multiple tissue types.

Sample size strongly influences accuracy (Fig. 2c). Tissues with more than 50 samples generally achieve greater than 90% accuracy, while those with fewer than 10 samples show variable performance. This relationship suggests that expanding training data for underrepresented tissues would further improve classification.

### Confusion patterns reflect biological relationships

Examination of the confusion matrix reveals that model errors are not random but instead reflect genuine biological relationships between tissues (Fig. 2b, Fig. 3). We identified three major confusion categories corresponding to distinct biological phenomena.

**Figure 3:**
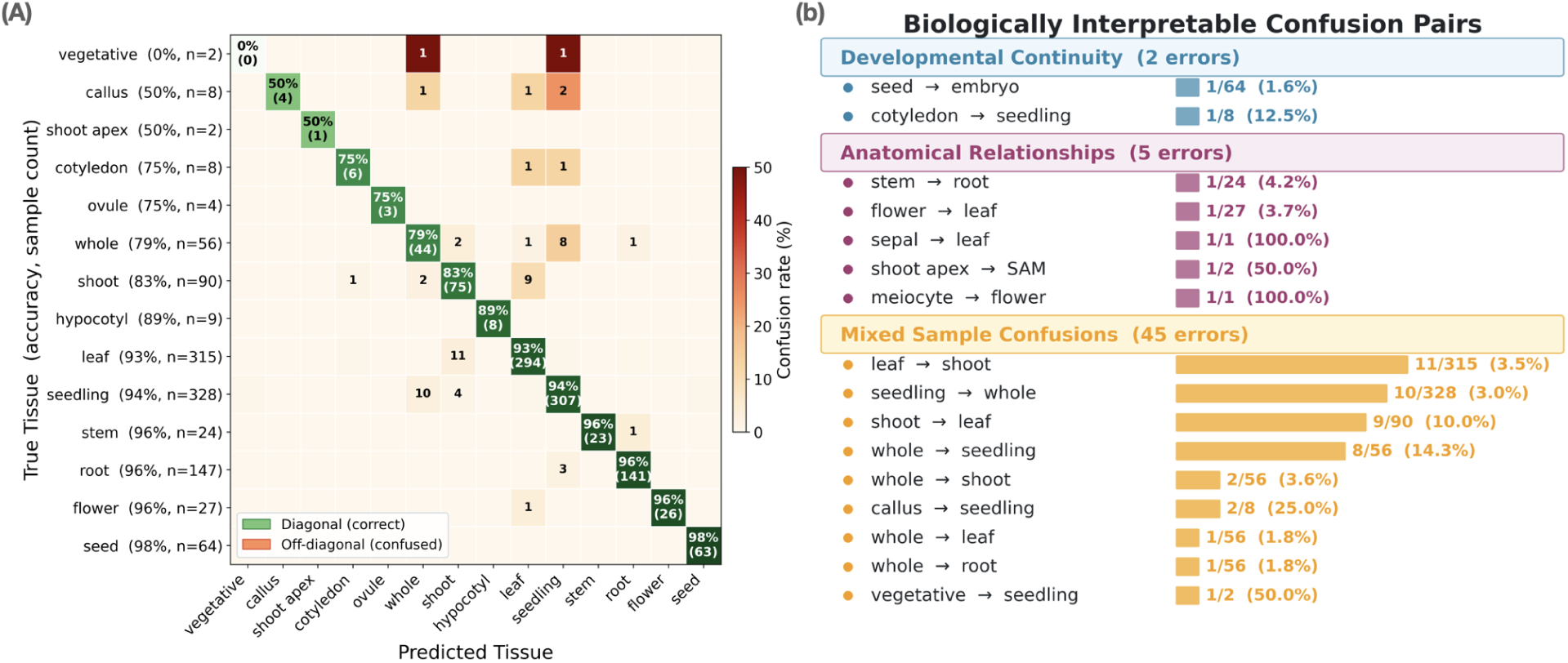
Confusion patterns in tissue classification. **(a)** Focused confusion heatmap showing 14 tissue types with < 100% classification accuracy on the validation set, sorted by accuracy (worst at top: vegetative at 0%, best at bottom: seed at 98%). Values are row-normalized to show the percentage of true tissue samples predicted as each class. Diagonal cells (green) display correct classification rate and count; off-diagonal cells (orange-red) display misclassification counts for confusion rates >= 1%. Tissues with 100% accuracy (e.g., pollen, fruit, inflorescence, ovule) are excluded for clarity. **(b)** Biologically interpretable confusion pairs grouped into three categories: developmental continuity (blue, 2 errors), anatomical relationships (magenta, 5 errors), and mixed sample confusions (amber, 45 errors). Each bar shows the misclassification count and percentage of the true tissue class. Biological explanations describe the mechanistic basis for each confusion pattern.

First, developmental continuity confusions occur between tissues at different stages of the same developmental program. Embryo samples are occasionally classified as seed (their containing structure), while cotyledon samples are classified as seedling (their developmental successor). These confusions reflect shared transcriptional programs across developmental transitions.

Second, anatomical relationship confusions occur between tissues that are physically related. Anther samples are sometimes classified as flower (which contains anthers), and sepal samples as leaf (to which sepals are morphologically similar). These patterns indicate that anatomically related tissues share expression signatures.

Third, mixed sample confusions involve samples representing multiple tissues. Whole plant and seedling samples show bidirectional confusion, as both contain mixtures of root, shoot, and leaf tissues. Similarly, “none” samples (unlabeled tissue) frequently match seedling expression patterns.

These biologically interpretable confusion patterns suggest that the model has learned genuine gene expression relationships rather than arbitrary classification rules.

### Attention importance reveals tissue-differential gene focus

To understand which genes drive classification decisions, we leveraged the self-attention mechanism of the transformer architecture. In each attention layer, every gene token computes attention weights over all other tokens, quantifying how much information it draws from each neighbor. These pairwise weights, averaged across 12 independently learned attention heads, provide two complementary views: (1) per-gene importance scores, obtained by summing how much attention each gene receives, and (2) pairwise gene-gene attention matrices, capturing which gene pairs the model processes jointly. We extract these from different parts of the architecture depending on the analysis goal (see Methods).

For tissue-differential gene importance analysis, we analyzed attention patterns from the shared encoder (Fig. 4a), which processes gene tokens before the architecture splits into tissue classification and MLM branches. For each sample, we extracted attention weights across all heads and layers, then aggregated them per gene to obtain importance scores. Grouping these scores by tissue reveals which genes the model considers most informative for each tissue type.

**Figure 4.**
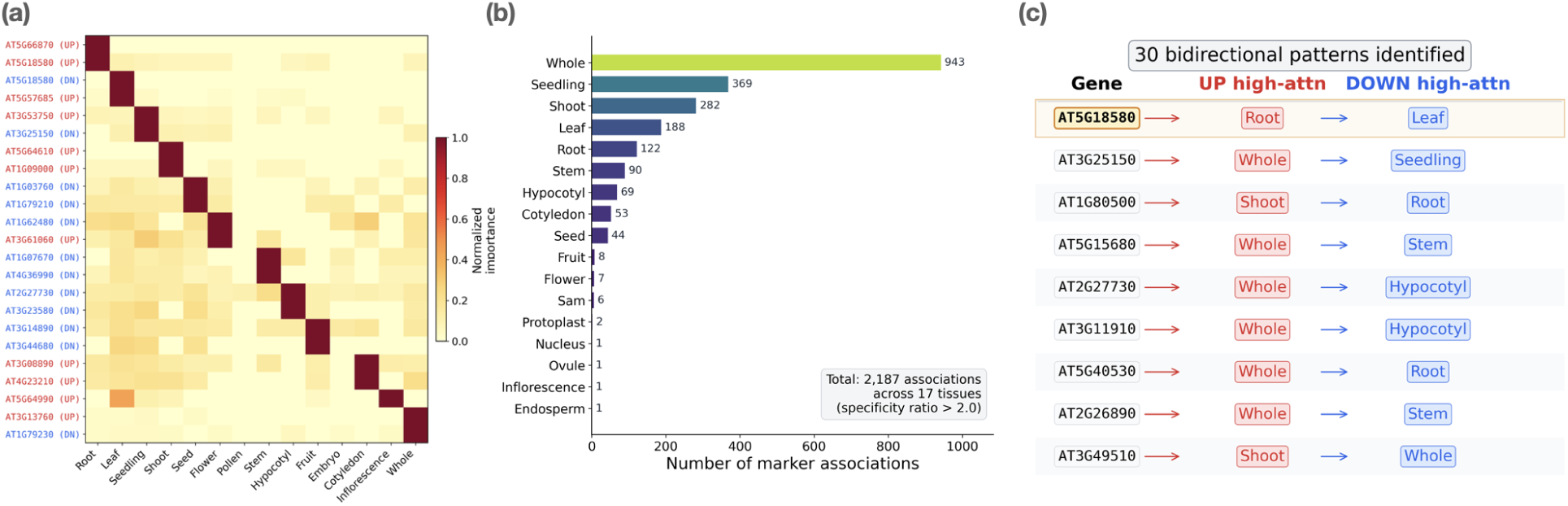
Attention importance reveals tissue-differential gene focus. **(a)** Heatmap of normalized gene importance scores for the top two high-attention genes per tissue across 15 major tissue types. Values are row-normalized (0-1 scale) to highlight tissue specificity. Gene labels are color-coded by expression direction: red for UP-regulated, blue for DOWN-regulated. **(b)** Distribution of high-attention gene associations across tissues. High-attention genes were defined as gene-tissue pairs with specificity ratio > 2.0 (importance in target tissue divided by mean importance across all tissues). A total of 2,187 associations were retained across 17 tissues with >50% classification accuracy. **(c)** Examples of bidirectional attention patterns --genes receiving high attention with opposite expression directions in different tissues (30 such genes identified). The highlighted example AT5G18580 receives high attention as UP in root (high expression is informative for root identity) and as DOWN in leaf (low expression is informative for leaf identity).

We computed specificity ratios as the importance of each gene for a target tissue divided by its mean importance across all tissues. Genes with specificity ratio greater than 2.0 were designated as high-attention genes—genes receiving disproportionately high attention in a given tissue context. This analysis identified 3,524 gene-tissue high-attention associations across the 47 tissue types, where the same gene can be flagged for multiple tissues with different expression directions.

To focus on reliable associations, we restricted the visualization in Fig. 4 to the 23 tissue types achieving greater than 50% classification accuracy (see Fig. 2), excluding tissues with insufficient samples or ambiguous expression signatures. This filtering retained 2,187 high-attention associations across 17 tissues (Fig. 4b; Table S4), representing the subset for which attention-derived importance scores are most reliable.

The distribution of high-attention genes across tissues is non-uniform, reflecting varying transcriptional distinctiveness (Fig. 4b). The whole plant leads with 943 associations—likely reflecting its mixed-tissue nature spanning multiple expression programs—followed by seedling (369) and shoot (282). Tissues with highly distinct transcriptomes such as root (122) have fewer total high-attention genes but much higher individual specificity ratios.

The heatmap of gene importance across tissues (Fig. 4a) displays the top 2 high-attention genes per tissue, with each row normalized by its maximum value across tissues to highlight tissue specificity. This row normalization means that for each gene, the tissue where it is most important appears as 1.0 (dark), while all other tissues are scaled relative to that maximum. A notable consequence is that pollen shows uniformly low values across all displayed genes. This is because pollen has zero individual high-attention genes—no single gene achieves a specificity ratio above 2.0 for pollen—and the genes selected as high-attention for other tissues (e.g., root, leaf) have near-zero importance in pollen. After row normalization, the pollen column is effectively blank. This does not indicate poor classification; pollen achieves 100% accuracy through combinatorial patterns of many co-occurring genes rather than any individual high-attention gene (see Figure S2).

Importantly, the same gene can receive high attention in different tissues with opposite directions (Fig. 4c). For example, AT5G18580.1_UP receives high attention in root (indicating high expression is informative), while AT5G18580.1_DOWN receives high attention in leaf (indicating low expression is informative). This bidirectional pattern reflects genuine tissue-specific regulation captured by the EES representation.

However, individual high-attention genes are not themselves tissue-specific in occurrence. AT5G18580.1_UP, for instance, has 10.5x attention specificity for root but actually appears in 27 of 47 tissues with only a 7.6% occurrence rate in root samples. This pattern is general: genes that the model attends to strongly for a given tissue are broadly distributed across the transcriptome, indicating that attention importance reflects the model’s classification strategy rather than restricted expression.

Notably, pollen achieves 100% classification accuracy despite having zero individual high-attention genes—no single gene exceeds the specificity threshold of 2.0 for pollen. Instead, pollen classification relies on the combinatorial co-occurrence of thousands of consistently silenced genes (see Figure S2), a fundamentally different strategy that foreshadows a key finding: tissue identity is encoded in gene combinations, not individual genes.

The gene importance analysis above used the shared encoder—the first four transformer layers that process gene tokens before the architecture splits into task-specific branches. This provides robust per-gene importance scores but cannot reveal how the model integrates these signals into a tissue decision. For that, we turn to the tissue classification branch, which adds two further layers of processing (six total) specifically trained for tissue discrimination. The tissue branch produces both sample-level embeddings (aggregate representations used for classification) and pairwise gene-gene attention patterns (which genes the model processes jointly when discriminating tissues).

### Learned embeddings capture tissue relationships

Each gene token in a sample passes through all four shared encoder layers and then through both tissue classification layers—six transformer layers in total. At the output of this pipeline, each gene token carries a 768-dimensional hidden state that has been progressively refined through self-attention interactions with all other genes. Mean-pooling these per-token hidden states across all gene positions produces a single 768-dimensional sample embedding (Fig. 5). This is the same representation that feeds the linear classification head: the model’s integrated summary of the sample’s gene expression pattern, compressed from a variable-length gene sequence into a fixed-size vector.

**Figure 5.**
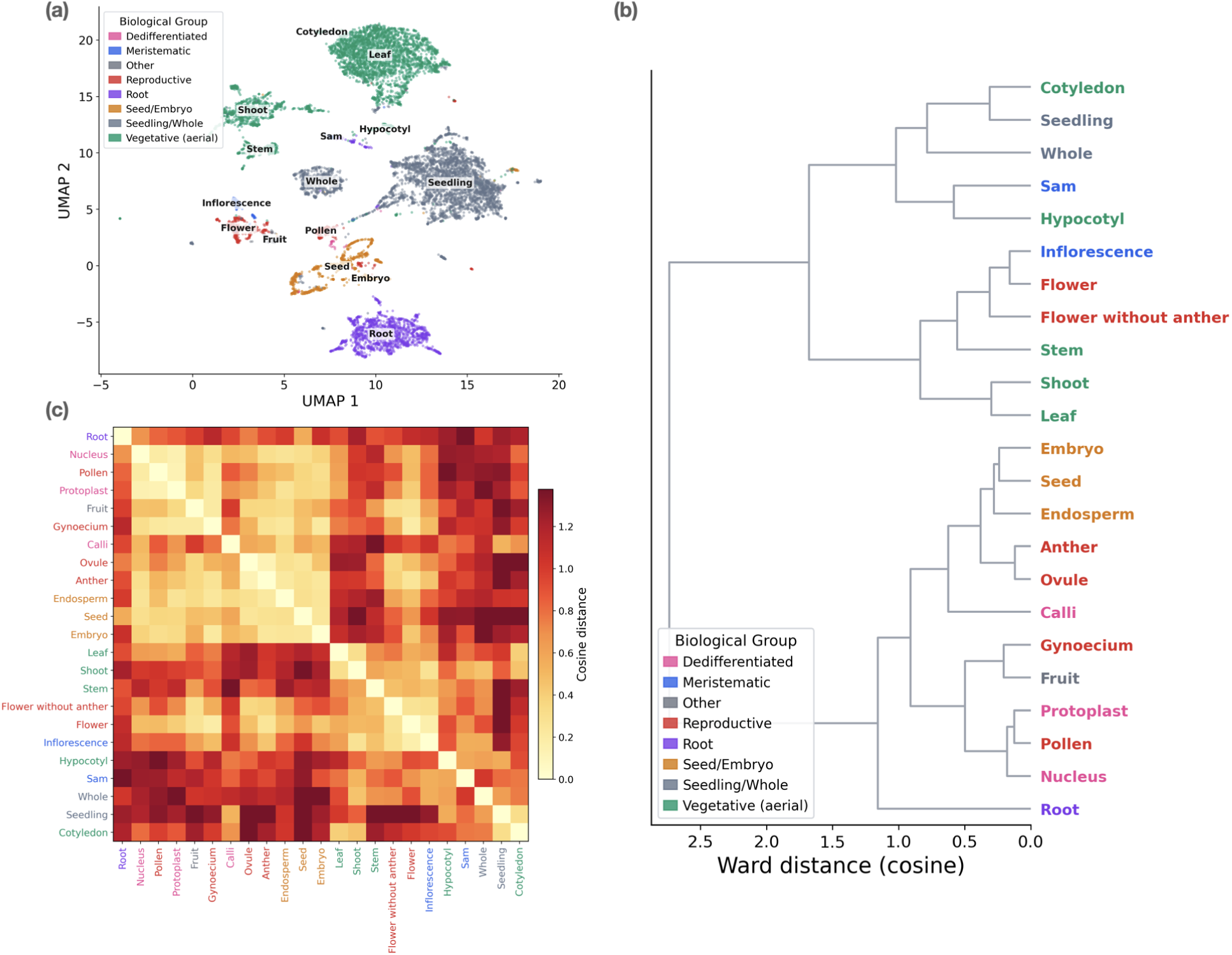
UMAP visualization and hierarchical analysis of learned tissue embeddings. **(a)** UMAP projection of 768-dimensional pooled representations from the tissue classification branch for all samples. Points are colored by biological groups: reproductive tissues (red), seed/embryo (amber), vegetative aerial tissues (green), root (violet), meristematic tissues (blue), seedling/whole plant (slate), and dedifferentiated tissues (pink). Tissue names are annotated for major clusters (n >= 30 samples). **(b)** Hierarchical clustering (Ward’s method) of tissue centroids computed as mean embeddings per tissue type, using cosine distance. Dendrogram leaf labels are colored by biological groups. **(c)** Heatmap of pairwise cosine distances between tissue centroids, ordered by the hierarchical clustering from panel (b). Lower values (lighter colors) indicate more similar tissue representations.

As with the gene importance analysis, we restricted the embedding visualization to the 23 tissue types with greater than 50% classification accuracy. UMAP visualization of these embeddings reveals clear tissue clustering, with reproductive tissues (pollen, flower, seed) separating from vegetative tissues (root, leaf, shoot) and developmental samples (seedling, embryo) forming intermediate groups (Fig. 5a).

Hierarchical clustering of tissue centroids (mean embedding per tissue) recapitulates expected biological groupings (Fig. 5b). Photosynthetic tissues cluster together, as do reproductive structures and seed-related tissues. This organization emerges entirely from learning to classify tissues from gene expression, without any explicit encoding of tissue relationships.

These embeddings enable potential transfer learning applications, where representations trained on tissue classification could initialize models for related tasks such as stress response prediction or developmental stage classification.

### Gene pairs are tissue-specific where individual genes are not

The gene importance analysis showed that individual high-attention genes are not tissue-specific in occurrence (e.g., AT5G18580.1_UP appears in 27 of 47 tissues despite 10.5x attention specificity for root). We asked whether gene *pairs* from the tissue classification branch show greater tissue specificity.

The tissue branch’s pairwise attention matrices—unlike the per-gene importance scores derived from the shared encoder—capture which gene pairs the model processes jointly when discriminating tissues. We extracted attention from the last layer of the tissue classification branch (layer six in the overall pipeline), where pairwise signals are sharpest; averaging across layers would blur these edges. We computed pairwise attention scores between all gene pairs, averaging across the 12 attention heads and tracking co-occurrence counts across all samples per tissue. To select reliable, high-attention edges, we retained only gene pairs with co-occurrence count >= 10 and attention weight above the tissue-specific median, then ranked by attention weight and selected the top 5,000 edges per tissue for analysis. For these high-confidence gene pairs, we then computed co-occurrence rates across the full 12,212-sample dataset: for a given gene pair, how often do both genes appear at expression extremes in the same sample, and in which tissue?

We quantified tissue specificity at two levels: for each gene pair (A, B), the *pair target rate* is the fraction of samples containing both A and B that belong to the target tissue; for comparison, each gene’s *single-gene target rate* is the fraction of all samples containing that gene that belong to the target tissue, and the single-gene baseline for an edge is the average of the two constituent genes’ rates (see Methods).

The contrast is striking (Fig. 6). Pollen shows the largest relative gain over individual genes: GRN gene pairs reach a mean pair target rate of 47.4% compared to a mean single-gene baseline of 17.4% (2.7-fold enrichment), reflecting pollen’s highly consistent transcriptional program despite comprising only ∼1% of total samples. Root pairs reach 53.2% vs 35.9% for the single-gene baseline (1.5-fold enrichment), with 26.4% of pairs exceeding 80% tissue specificity. Leaf pairs show a similar enrichment (47.8% vs 32.7%, 1.5-fold), consistent with leaf’s broader transcriptional overlap with other photosynthetic tissues.

**Figure 6.**
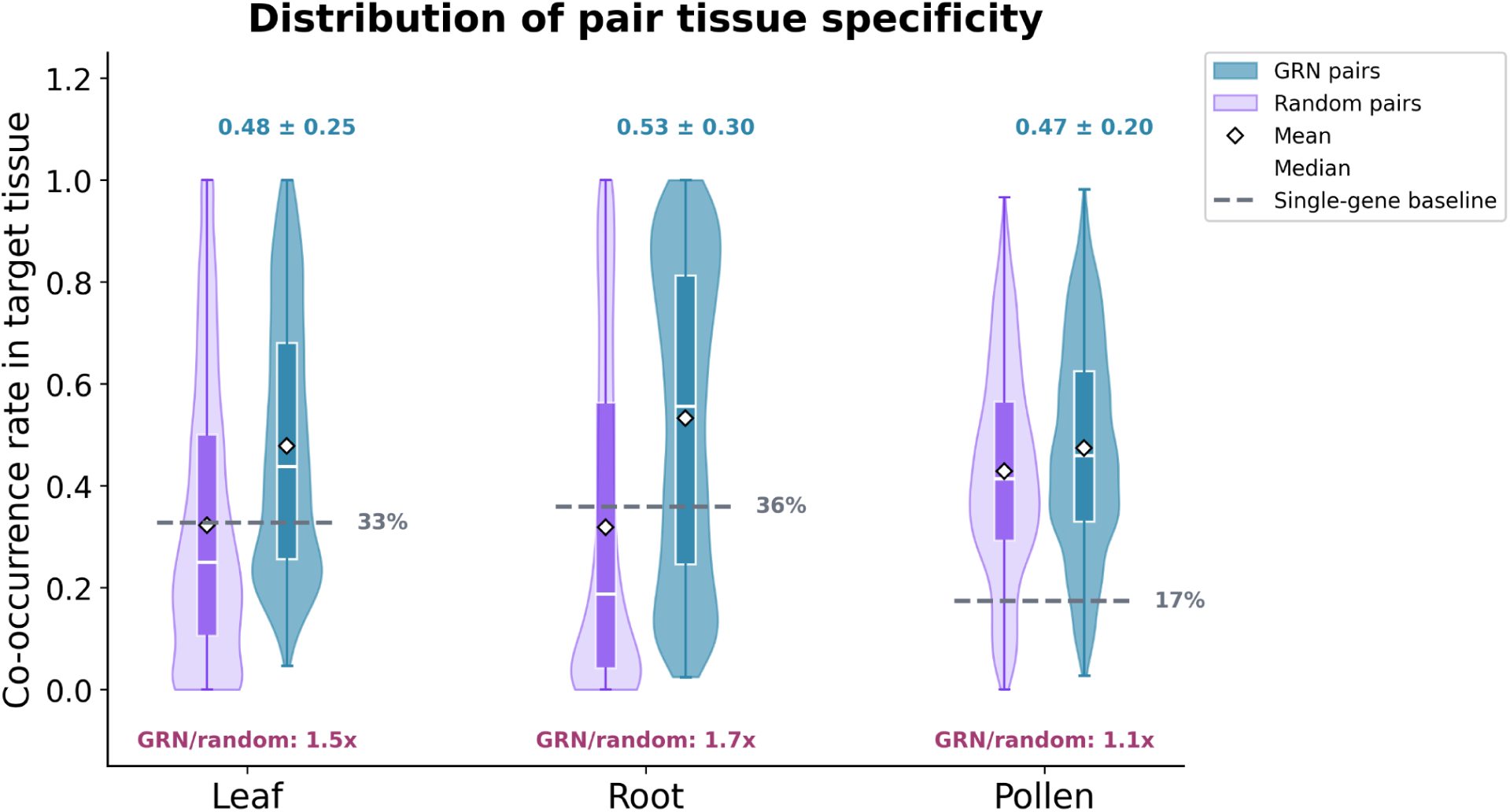
Gene pairs from attention-derived regulatory networks are more tissue-specific than individual genes. Violin plots show the full distribution of co-occurrence rates in the target tissue (i.e., the fraction of a gene pair’s total co-occurrences that fall in the target tissue) for 5,000 GRN gene pairs (blue) and 5,000 randomly paired genes from the same gene pool (purple) per tissue. Embedded mini-boxplots show quartiles; white diamonds indicate means. Dashed gray lines mark the single-gene baseline (mean fraction of individual gene occurrences in the target tissue). Mean ± s.d. annotations (blue) summarize each GRN distribution. GRN-to-random fold enrichments (magenta) quantify specificity gain beyond the trivial co-occurrence effect.

However, one might expect that any two genes co-occurring in the same sample would show higher tissue specificity than individual genes simply because co-occurrence is a more restrictive filter. To control for this effect, we constructed a random-pair null model: for each tissue, 5,000 gene pairs were randomly drawn from the same GRN gene pool (without regard to attention weights) and their co-occurrence specificity computed identically. Random pairs do show elevated specificity relative to individual genes (leaf: 32.3%, root: 31.8%, pollen: 42.8%), confirming the trivial pairing effect. Critically, however, GRN pairs remain substantially more specific than random pairs: 1.5-fold for leaf, 1.7-fold for root, and 1.1-fold for pollen (Fig. 6). The leaf and root enrichments demonstrate that the attention mechanism captures genuine co-regulatory relationships beyond what random pairing achieves. Pollen’s modest 1.1-fold enrichment is expected: because nearly all genes in pollen samples are co-silenced vegetative genes, even randomly paired genes show high co-occurrence specificity, leaving little room for GRN edges to improve.

The violin plot distributions reveal tissue-dependent patterns in distribution shape (Fig. 6). Root GRN pairs show a right-skewed distribution (s.d. = 0.30) with a substantial density at high specificity values, indicating many pairs with near-exclusive root occurrence. Pollen GRN pairs form a tight, symmetric peak around 0.47 (s.d. = 0.20). Leaf pairs peak at lower specificity values (∼0.25) with a long right tail (s.d. = 0.25), consistent with leaf’s transcriptional overlap with other photosynthetic tissues, diluting pair specificity for most edges while a subset of pairs achieves high tissue specificity. In all three tissues, the GRN pair distributions (blue violins) are shifted above both the random pair distributions (purple violins) and the single-gene baselines (dashed lines), confirming that the attention-derived combinatorial patterns capture tissue-specific co-regulation beyond what trivial gene pairing produces, consistent with the transformer’s attention mechanism.

This finding has important implications for how we interpret the model’s classification strategy. The transformer does not rely on individual genes with tissue-restricted expression. Instead, tissue identity is encoded in combinatorial gene expression patterns—specific pairs of genes at expression extremes—that are genuinely tissue-specific even when the individual genes are not. This explains why pollen achieves 100% classification accuracy with zero individual high-attention genes: its identity is defined by the coordinated silencing of thousands of gene pairs, a combinatorial signature that no single gene captures. The high tissue specificity of these pairwise attention patterns motivates constructing full gene regulatory networks from the attention weights.

### Attention-derived gene regulatory networks reveal tissue-specific programs

Given the tissue specificity of gene pairs identified above, we assembled comprehensive gene regulatory networks (GRNs) from the pairwise attention data. Edges observed in fewer than ten samples were filtered to retain only high-confidence interactions (Fig. 7).

**Figure 7.**
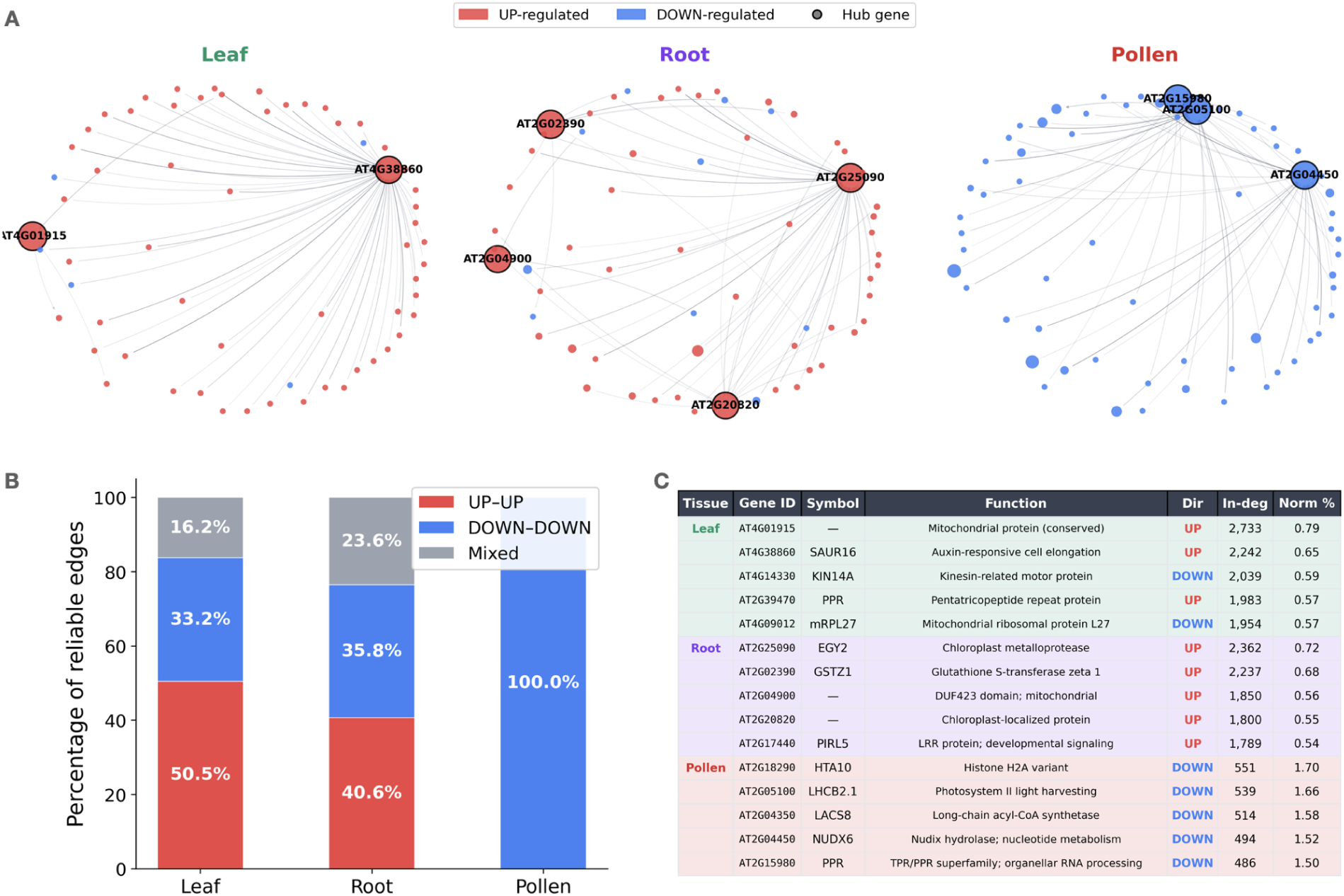
Tissue-specific gene regulatory networks (GRNs). **(a)** Ego-networks around the top five hub genes per tissue. Node color indicates expression direction (red = UP-regulated, blue = DOWN-regulated). Node size is proportional to attention in-degree. Hub genes are labeled and outlined in black. Edge opacity reflects attention weight. Only edges with co-occurrence count >= 10 and attention >= median are shown. **(b)** Edge directionality of reliable edges (count >= 10). Leaf and root show mixed directionality, while pollen consists entirely of DOWN-DOWN interactions, reflecting massive transcriptional silencing. **(c)** Hub gene annotations from TAIR/Araport11, showing gene symbol, biological function, expression direction, and attention in-degree.

Network statistics reveal tissue-dependent connectivity patterns. Using all available samples per tissue, leaf (3,004 samples) yielded 690,528 reliable edges (count >= 10) among 36,189,021 total gene pairs. Root (1,465 samples) produced 658,536 reliable edges from 13,477,411 pairs, while pollen (133 samples) produced 64,936 reliable edges from only 203,129 pairs—reflecting its specialized transcriptome. The degree distributions of all three networks exhibit heavy-tailed, scale-free-like topology with power-law exponents in the typical biological range (α = 2.17–2.98; Figure S3), consistent with known properties of biological networks [6].

A striking pattern emerges in edge directionality across tissues (Fig. 7b). Leaf GRN shows predominantly co-activation: 50.5% UP-UP, 33.2% DOWN-DOWN, with 16.2% mixed-direction edges. Root GRN shows 40.6% UP-UP, 35.8% DOWN-DOWN, and 23.6% mixed-direction edges—the highest proportion of cross-direction interactions, possibly reflecting the greater regulatory complexity of root tissue. In contrast, pollen GRN consists entirely of DOWN-DOWN interactions (100%), reflecting the massive transcriptional silencing of vegetative genes characteristic of this specialized reproductive cell type.

Hub gene analysis with biological annotation reveals tissue-specific regulatory programs (Fig. 7c). Leaf hub genes include both up-regulated and down-regulated genes spanning diverse functions: an uncharacterized protein (AT4G01915; 2,733 incoming edges), a SAUR-like auxin-responsive protein (AT4G38860; 2,242 edges), and a P-loop NTPase superfamily protein (AT4G14330; 2,039 edges). Additional leaf hubs include PnsL1/PsbP-like protein 2 (AT2G39470; photosystem component; 1,983 edges) and mitochondrial ribosomal protein L27 (AT4G09012; 1,954 edges), linking leaf identity to photosynthetic function and organellar protein homeostasis.

Root hub genes are predominantly up-regulated: CIPK16 (AT2G25090; CBL-interacting protein kinase; 2,362 edges), GSTZ1 (AT2G02390; glutathione S-transferase zeta 1; 2,237 edges), and PIRL5 (AT2G17440; plant intracellular ras group-related LRR protein; 1,789 edges). The prevalence of up-regulated stress signaling and detoxification genes among root hubs reflects the root’s active metabolic program and its role in environmental sensing.

Pollen hub genes are exclusively down-regulated, spanning diverse vegetative functions: APC10 (AT2G18290; anaphase promoting complex 10; 551 edges), LHCB2.1 (AT2G05100; photosystem II light harvesting; 539 edges), LACS8 (AT2G04350; AMP-dependent synthetase and ligase; 514 edges), and NUDT6 (AT2G04450; nudix hydrolase; 494 edges). This diversity—spanning cell cycle regulation, photosynthesis, lipid metabolism, and nucleotide metabolism—reflects the breadth of vegetative programs coordinately silenced in pollen.

Although pollen hub genes have lower raw in-degree counts than leaf or root hubs, normalizing by total high-attention edges reveals that pollen hubs are proportionally the most connected: the top pollen hub (APC10) accounts for 1.70% of all high-attention edges, compared to 0.79% for the top leaf hub and 0.72% for root (Fig. 7c). This reflects pollen’s smaller but more focused network, where attention is concentrated on a compact set of coordinately silenced genes.

### Token co-occurrence determines GRN reliability

To understand why pollen GRN shows such consistent patterns, we analyzed token co-occurrence across 100 randomly sampled samples per tissue (Table S5). Token diversity varies dramatically between tissues: leaf samples collectively use 38,517 unique tokens with an average reuse rate of 4.2 times per token, root samples use 37,329 unique tokens with a reuse rate of 8.8 times, while pollen samples use 27,030 unique tokens but with a 12-fold higher reuse rate than leaf (49.2 times per token).

Critically, in pollen 1,211 tokens appear in all 100 sampled samples (100% frequency), of which 1,182 are DOWN-regulated and only 29 are UP-regulated; an additional 9,685 tokens exceed 90% frequency. No tokens reach 100% frequency in leaf or root. This explains the clarity of pollen GRN—the same genes are consistently silenced across all pollen samples, creating highly reliable attention patterns. In contrast, leaf and root samples show greater token diversity, reflecting their cellular heterogeneity as bulk tissue samples containing multiple cell types.

Comparing the top 500 edges across tissues reveals complete specificity: leaf and root share zero edges, leaf and pollen share zero edges, and root and pollen share zero edges among their strongest connections. This complete lack of overlap demonstrates that the model has learned genuinely tissue-specific regulatory patterns rather than generic co-expression relationships.

## Discussion

We have presented EES-Transformer V2, a dual-path transformer architecture for tissue classification and gene representation learning from RNA-seq data. The architecture physically separates classification and generative objectives, preventing information leakage between tasks. Applied to *Arabidopsis thaliana*, the model achieves 91-92% accuracy across 47 tissue types while providing interpretable attention-based gene importance analysis and tissue-specific regulatory network inference.

The Extreme Expression Set representation offers a principled approach to discretizing continuous expression data. By focusing on genes at expression extremes, EES captures the most tissue-informative signals while reducing computational requirements. The success of this representation suggests that tissue identity is indeed encoded primarily in exceptionally expressed genes—a hypothesis with implications for how we conceptualize tissue-specific transcriptional programs.

The dual-path architecture ensures fair evaluation of tissue classification by preventing information leakage from labels to predictions. The 91-92% accuracy demonstrates that EES patterns alone contain sufficient information for tissue identification. The separation also enables meaningful interpretation of attention patterns, as the classification branch must rely entirely on learned gene relationships.

A notable consequence of the multi-task design is that the 15% random masking required for MLM also applies to the tissue classification branch, since both branches share the same masked input. This stochastic masking, combined with random token order shuffling at each access, functions as token-level dropout for classification: the model must identify tissue identity from a different ∼85% subset of genes every time. This regularization effect likely contributes to the model’s generalization, as the classifier cannot memorize specific gene combinations but must instead learn distributed tissue signatures across the full gene set. It also means that classification accuracy without masking would likely be higher than the reported 91-92%.

The biologically meaningful confusion patterns provide confidence that the model has learned genuine biology rather than dataset artifacts. Confusions between developmentally related tissues (embryo-seed, cotyledon-seedling), anatomically related tissues (anther-flower, sepal-leaf), and mixed samples (whole-seedling) all correspond to known biological relationships. This interpretability distinguishes our approach from black-box classifiers and suggests applications in quality control, where unexpected classifications could flag annotation errors.

The attention-based analysis reveals an important distinction between individual gene importance and combinatorial specificity. While individual high-attention genes reflect the model’s classification strategy, they are not tissue-specific in occurrence—a gene receiving 10x attention specificity for root may appear in 27 of 47 tissues. In contrast, gene pairs from attention-derived regulatory networks show higher tissue specificity, with pollen pairs showing 2.7-fold enrichment and root and leaf pairs each showing 1.5-fold enrichment over single-gene baselines. This finding supports the interpretation that transformers capture combinatorial patterns rather than relying on individual features, consistent with the pollen observation where classification succeeds through coordinated silencing of gene pairs despite zero individual high-attention genes. The identification of bidirectional attention patterns—genes whose high expression is informative in one tissue while low expression is informative in another—further demonstrates that the EES representation captures regulatory relationships beyond simple on/off patterns.

The extension of attention analysis to gene regulatory network inference reveals tissue-specific regulatory structures with biological meaning (Fig. 7). Hub gene annotation against Ensembl Plants/Araport11 demonstrates that the model recovers biologically coherent programs without supervision (Table S7). Leaf hubs include auxin-responsive proteins, a photosystem component (PnsL1/PsbP-like protein 2), and mitochondrial ribosomal protein L27, reflecting leaf-specific photosynthetic function and organellar protein homeostasis. Root hubs feature a

CBL-interacting protein kinase (CIPK16), glutathione S-transferase (GSTZ1), and a developmental LRR protein (PIRL5), consistent with the root’s roles in stress signaling and nutrient uptake, while several uncharacterized hub genes represent candidates for future functional studies. The complete dominance of DOWN-DOWN interactions in pollen GRN is particularly striking: pollen hub genes span photosynthesis (LHCB2.1), lipid metabolism (LACS8), nucleotide metabolism (NUDT6), cell cycle regulation (APC10), and chloroplast RNA polymerase (rpoB), reflecting the breadth of vegetative programs silenced in this specialized cell type. Notably, normalizing hub in-degree by each tissue’s total high-attention edges reveals that pollen hubs are proportionally the most connected (1.70% for the top hub, vs. 0.79% in leaf and 0.72% in root), despite having lower raw edge counts. This indicates that attention is more concentrated in pollen’s compact network, consistent with the coordinated silencing of a focused set of vegetative genes. The high token reuse rate in pollen (49-fold vs. 4-fold in leaf) further reflects this transcriptional specialization and produces highly reliable network structure. These findings suggest that attention-derived GRNs may be most informative for homogeneous cell populations, pointing toward applications in single-cell transcriptomics where each cluster represents a relatively pure cell type. More broadly, the EES representation strategy is well suited to single-cell RNA-seq data: individual cells of a given type share more consistent expression profiles than bulk tissue samples, which average over heterogeneous cell populations. The pollen results illustrate this principle—pollen samples behave like a near-homogeneous cell population, yielding the highest token reuse rate (49.2x) and the most focused GRN structure. Single-cell clusters would similarly exhibit high token consistency, enabling the EES framework to capture cell-type-specific extreme expression signatures with greater precision than is achievable from bulk data. Furthermore, the EES discretization is inherently scalable: because continuous expression values are reduced to binary extreme states (UP/DOWN), the representation remains compact regardless of dataset size. Single-cell atlases routinely comprise hundreds of thousands of cells across dozens to hundreds of cell types—a scale that would strengthen both the transformer’s tissue classification and the reliability of attention-derived GRNs, as more samples per cell type increase token co-occurrence counts and network confidence.

Importantly, attention weights reveal co-attention patterns rather than direct regulatory relationships. Recent methods such as STGRNS [7] and AttentionGRN [8] have similarly leveraged transformer attention for GRN inference, though these approaches focus on learning attention patterns specifically for regulatory relationship prediction rather than extracting post-hoc networks from classification models. High attention between genes A and B indicates that the model considers B’s representation informative when processing A—suggesting shared regulatory mechanisms or co-expression—but does not establish causation. Validation against experimentally determined regulatory networks (e.g., DAP-seq, ChIP-seq) or curated gene expression databases would be valuable to assess the biological validity of attention-derived edges.

Several limitations suggest directions for future work. First, the model is trained specifically on Arabidopsis and would require retraining for other species. Transfer learning approaches could potentially address this by learning species-invariant gene representations. Second, tissues with few samples show unreliable performance, indicating the need for data augmentation or few-shot learning techniques. Third, attention-based importance does not establish causal relationships between genes and tissue identity; experimental validation remains necessary to confirm the biological significance of high-attention genes and gene pairs.

Future directions include extending the approach to other model organisms and crops, integrating temporal dynamics through sequential expression data, and combining EES representations with other molecular data types such as chromatin accessibility. Applying EES-Transformer to single-cell RNA-seq data is a particularly promising direction: single-cell profiles from defined cell types would provide the consistent expression patterns that our pollen analysis demonstrates are ideal for the EES framework, and the large sample sizes typical of single-cell atlases (10^5–10^6 cells across hundreds of cell types) would directly strengthen both classification accuracy and GRN reliability by increasing token co-occurrence counts per cell type. The dual-path architecture is also applicable beyond tissue classification to any multi-task setting where fair evaluation of one task requires isolation from information available to another.

In summary, EES-Transformer V2 provides accurate tissue classification with interpretable attention-based gene importance analysis and attention-derived gene regulatory networks. The key biological insight—that tissue identity is encoded in combinatorial gene expression patterns rather than individual genes—emerges from comparing single-gene attention importance (broadly distributed) with gene pair co-occurrence (tissue-specific). Hub gene annotation reveals biologically coherent programs—photosynthetic and signaling genes in leaf, stress response kinases in root, and silenced vegetative genes in pollen—validating the approach as a framework for combining classification, representation learning, and regulatory network inference in transcriptomics.

## Methods

### Data collection and preprocessing

We compiled RNA-seq data from publicly available Arabidopsis thaliana studies, obtaining 12,212 samples annotated with 47 tissue types. Raw sequencing data were processed through a standardized pipeline: quality control with FastQC, adapter trimming with Trimmomatic, alignment to the TAIR10 reference genome with STAR, and expression quantification with RSEM. Expression values were normalized to transcripts per million (TPM) for cross-sample comparability.

For EES sentence construction, we first filtered genes with mean TPM below 1.0 across all samples to remove lowly expressed genes, retaining 30,647 genes. For each gene, we computed expression percentiles across all samples to establish gene-specific thresholds. For each sample, genes with TPM above the 95th percentile for that gene received an “_UP” suffix, while genes below the 5th percentile received “_DOWN”. The resulting tokens were concatenated into space-separated sentences, one per sample. The vocabulary comprises 46,728 tokens (30,647 UP tokens, 16,076 DOWN tokens, and 5 special tokens; Table S6), reflecting that not all genes appear at both expression extremes.

The choice of 95th and 5th percentile thresholds is motivated by three considerations. First, these thresholds correspond approximately to two standard deviations from the mean in a normal distribution, a standard practice for identifying differentially expressed genes. Second, selecting the top and bottom 5% of genes per sample (approximately 10% total) provides sufficient signal for tissue identification while maintaining computationally tractable sequence lengths (median 1,542 tokens per sample). Third, genes at expression extremes are more likely to reflect tissue-specific transcriptional programs, as constitutively expressed housekeeping genes typically fall within the middle percentiles.

### Model architecture

EES-Transformer V2 consists of three components: a shared encoder, a tissue classification branch, and a masked language modeling branch.

**Embedding layer.** Input tokens are embedded into 768-dimensional vectors through a learned embedding table. No positional encodings are added, as gene sets are permutation invariant.

**Shared encoder.** Four transformer layers process the embedded tokens. Each layer comprises multi-head self-attention (12 heads, 64 dimensions per head) followed by a position-wise feed-forward network (768 → 3072 → 768) with GELU activation. Layer normalization and residual connections follow standard transformer design.

**Tissue classification branch.** Two additional transformer layers (same configuration as shared encoder) process the shared representations. Mean pooling across all tokens produces a 768-dimensional sample representation, which is projected through a linear layer to 47 tissue class logits. This branch receives no tissue label information at any point.

**MLM branch.** Tissue labels are encoded through a learned embedding table (47 tissues × 768 dimensions). The tissue embedding is added to the shared representations at each token position. Four transformer layers process these tissue-conditioned representations, followed by a linear projection to vocabulary logits (46,728 dimensions).

The total model comprises approximately 144.6 million parameters (Table S3).

### Training procedure

The model was trained with a multi-task objective combining tissue classification and masked language modeling:

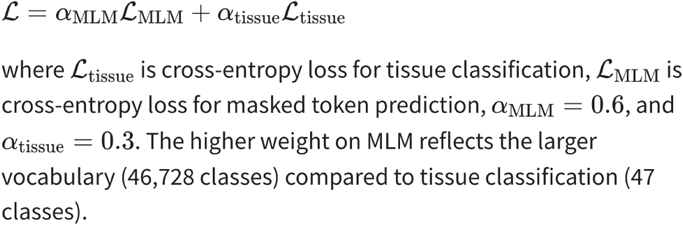

For MLM, we randomly mask 15% of input tokens, replacing 80% with a [MASK] token, 10% with a random token, and 10% unchanged, following BERT conventions [9]. Additionally, gene token order is randomly shuffled at each access, as EES sequences are permutation invariant.

Importantly, the masked input is shared by both branches: the same 15%-masked token sequence passes through the shared encoder and then feeds into both the tissue classification and MLM branches. Because the masking is stochastic—different random tokens are masked and the gene order is reshuffled every time a sample is accessed—the tissue classifier never sees the identical input twice for the same sample. This effectively serves as token-level dropout for the classification branch, preventing the model from relying on any individual gene and encouraging robust, distributed representations of tissue identity. The reported classification accuracy therefore reflects performance under this input noise regime.

We used the AdamW optimizer with learning rate 5e-5, weight decay 0.01, and linear warmup over 1,000 steps followed by cosine decay. Batch size was 2 due to the long sequence lengths (up to 2,048 tokens). Training proceeded for up to 100 epochs with early stopping based on validation tissue accuracy (patience = 10 epochs), converging at epoch 65.

Data were split 80%/10%/10% for training, validation, and testing, stratified by tissue type when possible. Maximum sequence length was set to 2,048 tokens; longer sequences were truncated and shorter sequences padded.

### Evaluation metrics

Tissue classification accuracy was computed as the fraction of correctly classified samples. We report both overall accuracy (macro-averaged across samples) and per-tissue accuracy (fraction correct within each tissue). Confidence was computed as the softmax probability assigned to the predicted class.

Confusion matrices were computed by counting predictions for each true-predicted tissue pair, normalized by row to show the distribution of predictions for each true tissue.

### Attention weight extraction

The EES-Transformer’s self-attention mechanism provides interpretable information about gene relationships. In each transformer layer, every gene token attends to all other tokens through 12 independent attention heads (64 dimensions each), producing attention weights that quantify pairwise token interactions. We exploit these weights for two distinct analyses—gene importance analysis and gene regulatory network inference—using different parts of the architecture tailored to each objective.

### Attention-based gene importance analysis

To extract gene importance scores, we performed inference on all training and validation samples, extracting attention weights from the **shared encoder**. For each sample, we obtained attention matrices of shape (layers × heads × tokens × tokens) from the **four shared encoder layers**.

We used the shared encoder rather than the tissue-specific branch because the shared encoder processes gene tokens before the architecture splits into downstream tasks. Its attention reflects intrinsic gene co-expression relationships that are informative for tissue classification without being optimized solely for that objective. Importance scores were averaged across all four layers to provide robust per-gene estimates.

For each gene token in a sample, we computed its raw importance as the total attention weight it receives from all tokens (including self-attention), averaged across all attention heads and layers:

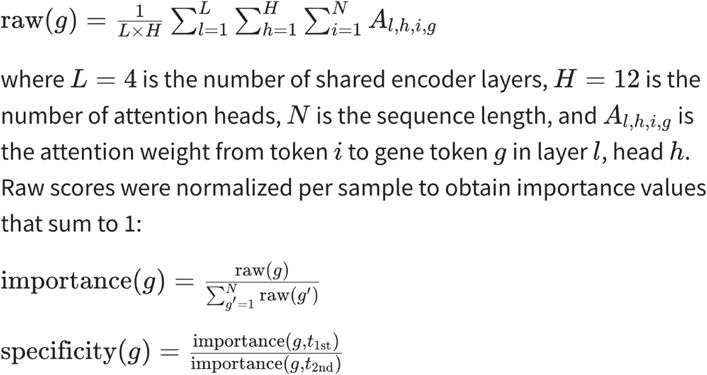

Gene importance scores were aggregated by tissue by averaging across all samples of each tissue type. The specificity ratio for gene *g* was computed as the ratio of its importance in its top-ranked tissue to its importance in the second-highest tissue:

Genes with specificity ratio > 2.0 were designated as high-attention genes. We use “high-attention gene” rather than “marker gene” because attention importance reflects the model’s classification strategy rather than tissue-specific expression; empirically, high-attention genes appear broadly across tissues.

### Attention-based gene regulatory network extraction

We extracted tissue-specific gene regulatory networks from attention weights using the following procedure:

**Sample selection.** We used all available samples for each tissue type (leaf: 3,004; root: 1,465; pollen: 133) to maximize statistical power and ensure reproducible network inference.

**Attention extraction.** For each sample, we performed inference and extracted the attention matrices from the tissue encoder (last layer). We used the tissue branch’s last layer rather than the shared encoder because GRN inference requires tissue-specific pairwise relationships. The last layer of the tissue branch sits at position 6 in the overall six-layer processing pipeline (four shared + two tissue-specific), where attention patterns are most refined for tissue discrimination. Using only the final layer preserves sharp pairwise signals, whereas averaging across layers would dilute edge specificity. Attention weights were averaged across all 12 attention heads to obtain a single attention matrix per sample:

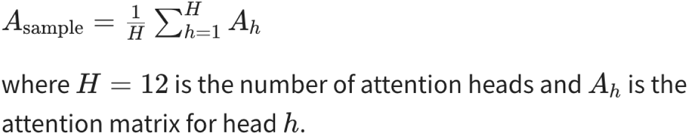

**Gene pair aggregation.** Unlike position-based attention analysis, we tracked attention by gene pair names (e.g., AT1G01010_UP → AT2G02020_DOWN) rather than positional indices, as gene order varies between samples. For each gene pair observed in a sample, we recorded the attention weight and incremented a co-occurrence counter.

**Network consolidation.** Across all samples for a tissue, we computed the mean attention weight for each gene pair and retained only edges with co-occurrence count >=10 to filter spurious interactions:

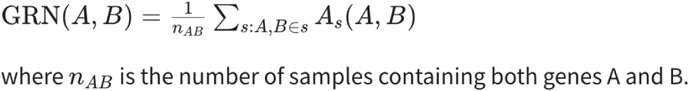

**Direction analysis.** Gene pairs were classified by their expression direction: UP→UP, DOWN→DOWN, UP→DOWN, or DOWN→UP, based on the token suffixes.

**Hub gene identification.** Hub genes were identified by computing the attention in-degree (number of incoming attention edges with weight above the median) for each gene. We use “attention in-degree” to distinguish from true regulatory in-degree, as attention weights reflect co-processing patterns rather than established regulatory relationships.

**Co-occurrence tissue specificity validation.** To validate whether GRN edges represent genuinely tissue-specific relationships, we computed co-occurrence tissue specificity for the top 5,000 edges per tissue. We selected edges with co-occurrence count >= 10 and attention weight above the median, ranked by attention weight, and retained the top 5,000 per tissue. For each selected gene pair (A, B), we identified all samples across the full 12,212-sample dataset where both genes appear at expression extremes. The *pair target rate* was computed as the fraction of co-occurrences occurring in the GRN’s source tissue. Because the two genes in a pair typically have different occurrence patterns, the single-gene baseline for each edge is the average of the two constituent genes’ rates. This per-edge baseline is then averaged across all 5,000 edges to produce the tissue-level single-gene mean reported in the Results.

### Software and hardware

The model was implemented in PyTorch 2.0 and trained in FP32 precision on a single NVIDIA GeForce RTX 4090 GPU (24GB). Training required approximately 19 hours to convergence (65 epochs with early stopping, best validation accuracy at epoch 63). Code is available at https://github.com/k821209/ees-transformer. Pre-trained model weights are available at https://huggingface.co/dirmy/ees-transformer-arabidopsis.

## Data Availability

The processed EES sentences, vocabulary files, and tissue annotations are available at https://github.com/k821209/ees-transformer. Raw RNA-seq data were obtained from public repositories (SRA accessions listed in Table S1).

## Code Availability

The EES-Transformer V2 code, including model architecture, training scripts, and analysis notebooks, is available at https://github.com/k821209/ees-transformer. Pre-trained model weights are available at https://huggingface.co/dirmy/ees-transformer-arabidopsis.

## Acknowledgements

Author Contributions

J.-S.P. developed the bioinformatics pipeline, implemented the model architecture, and performed all computational analyses. Y.L. collected and curated the RNA-seq dataset. Y.J.K. conceived and designed the study, and wrote the manuscript. All authors reviewed and approved the final manuscript.

## Competing Interests

The authors declare no competing interests.

## Notes

### Competing Interest Statement

The authors have declared no competing interest.

### Funding

- Government of the Republic of Korea, MSIT: RS-2024-00336161

