## SupplementaryFigures for "EES-Transformer: A Dual-Path Transformer for Tissue Classification and Gene Representation Learning from Extreme Expression Sets": SupplementaryMaterials-ees.docx

### Supplementary Materials

#### EES-Transformer: A Dual-Path Transformer Architecture for Tissue Classification and Gene Regulatory Network Inference from Arabidopsis RNA-seq Data

#### Supplementary Figures

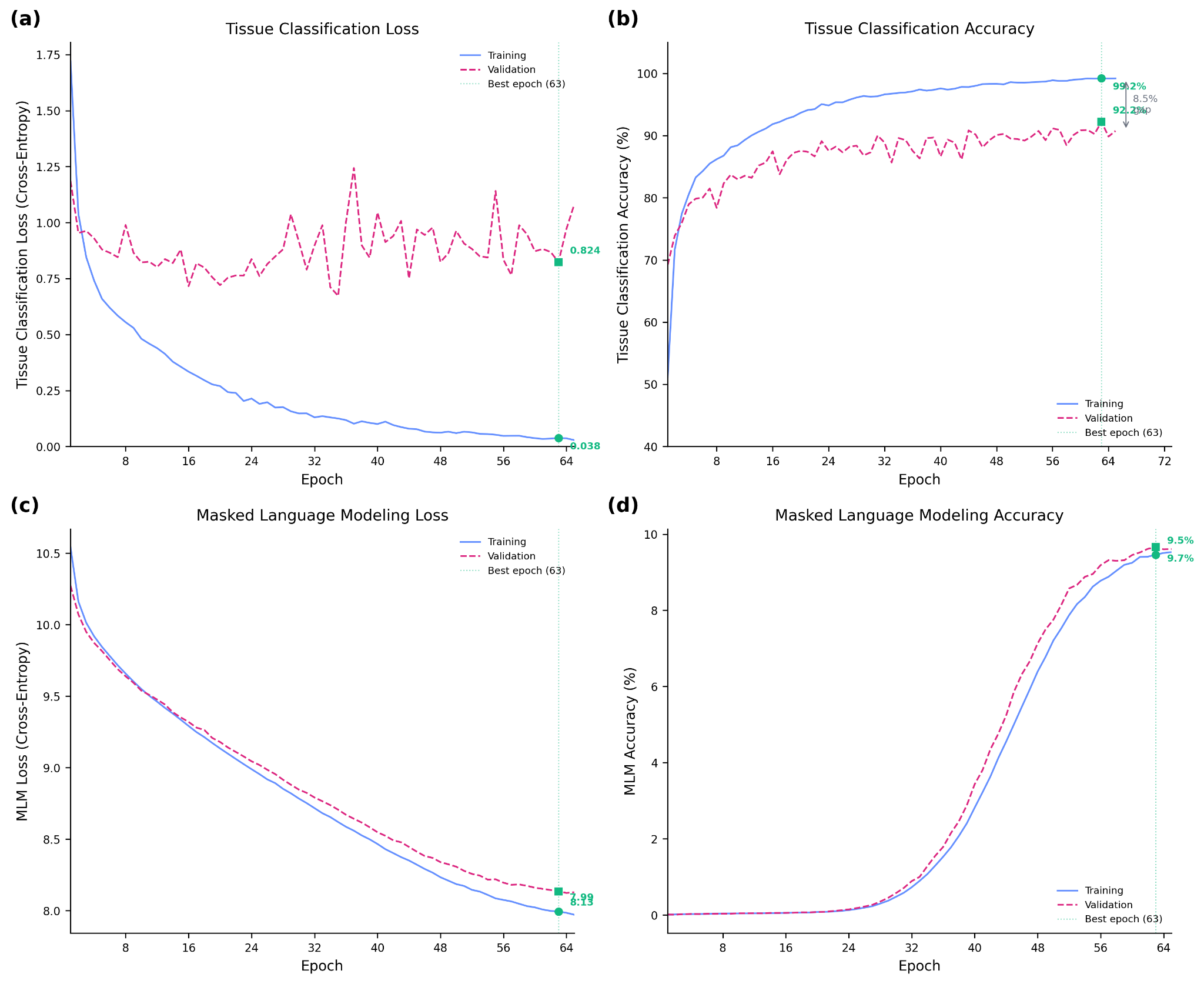

##### Figure S1: Training Convergence Curves

Training dynamics over 62 epochs for EES-Transformer V2. **(a)** Training and validation loss for tissue classification (cross-entropy). Tissue classification loss decreases rapidly in the first 10 epochs and stabilizes near 0.03 (training) and 0.9 (validation) by epoch 60. **(b)** Training and validation accuracy for tissue classification. Training accuracy reaches 99.2% while validation accuracy plateaus at 92.2%, with a ~7% gap indicating mild overfitting. **(c)** MLM loss tracks closely between training and validation sets throughout training, showing no overfitting on the masked language modeling task. **(d)** Combined training summary panel showing all metrics.

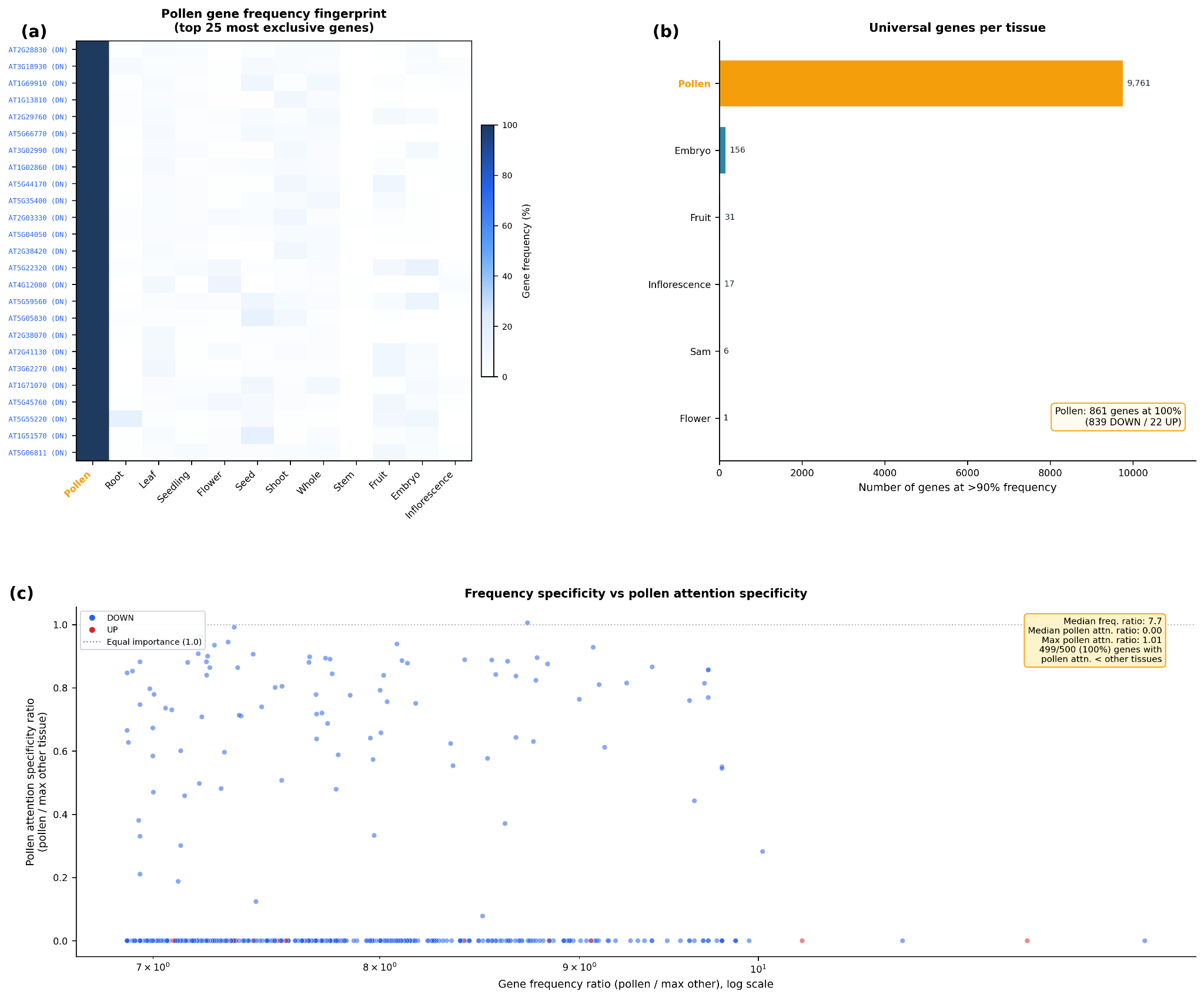

##### Figure S4: Pollen Combinatorial Marker Analysis

**(a)** Heatmap of gene frequency (% of samples containing each gene) for the 25 most pollen-exclusive genes across 12 major tissues. Pollen column is uniformly dark (100% frequency) while other tissues show near-zero frequency, demonstrating the distinctive pollen gene frequency fingerprint. Genes are ranked by frequency ratio (pollen / max other major tissue). **(b)** Number of "universal" genes (>90% frequency) per tissue (tissues with >=20 samples). Pollen dominates with 9,761 universal genes -- orders of magnitude more than any other tissue. Annotation: 861 genes at 100% frequency (839 DOWN / 22 UP). **(c)** Scatter plot comparing two pollen-specificity metrics for the top 500 pollen-enriched genes. X-axis: gene frequency ratio (pollen / max other tissue), measuring how exclusively a gene appears in pollen EES sentences. Y-axis: pollen attention specificity ratio (pollen importance / max other tissue importance), measuring how much more attention the model allocates to this gene when processing pollen versus any other tissue. Dotted gray line indicates ratio = 1.0 (equal importance). The disconnect is striking: genes with 7–12× pollen frequency specificity have pollen attention ratios clustered near zero (median = 0.00, max = 1.01), meaning the model gives them equal or less attention in pollen compared to other tissues. This demonstrates that pollen classification relies on the combinatorial co-occurrence of thousands of genes rather than elevated attention to any individual gene.

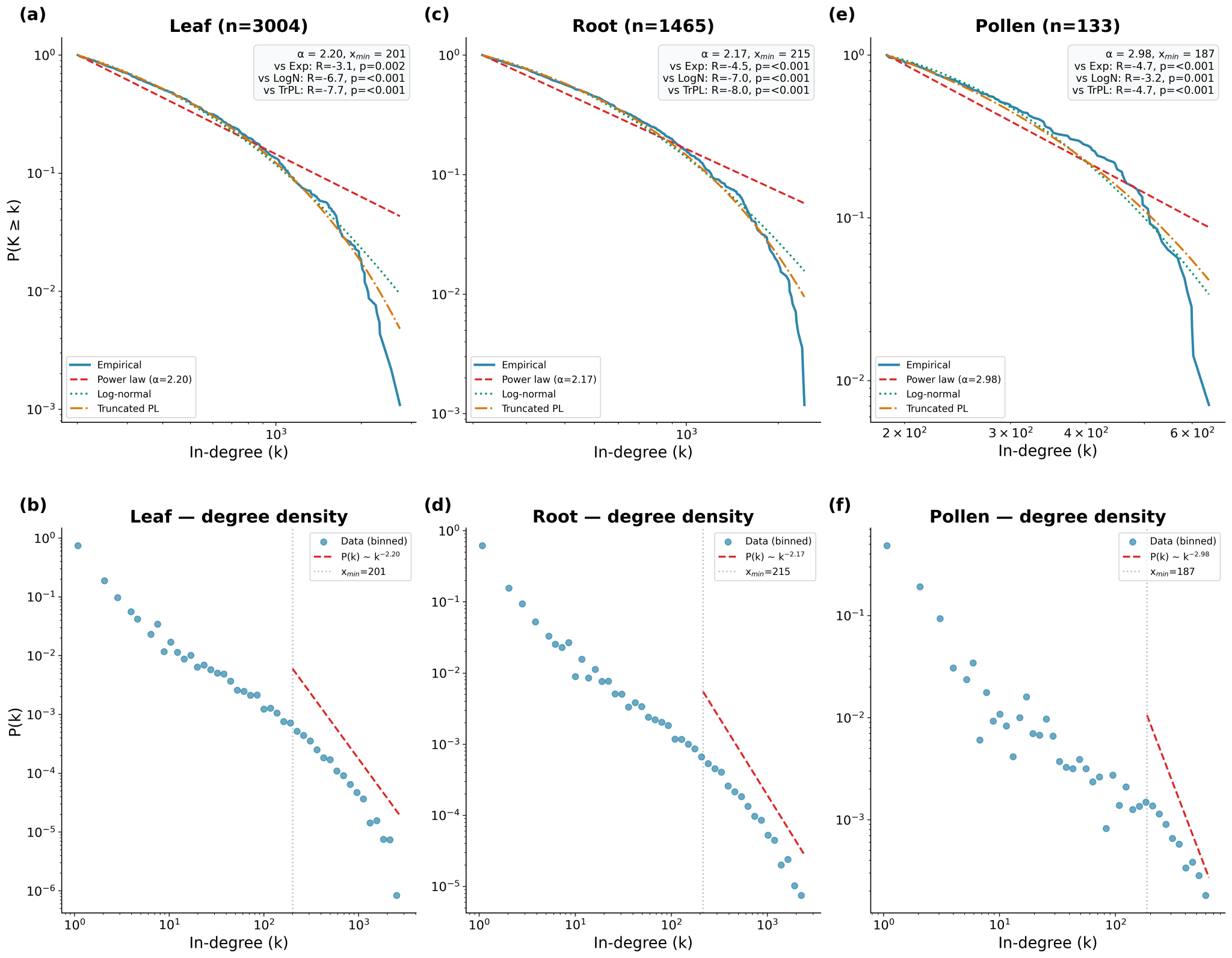

##### Figure S5: Scale-Free Network Analysis of Attention-Derived GRNs

**(a–c)** Complementary cumulative distribution function (CCDF) of in-degree for leaf, root, and pollen GRNs (edges with co-occurrence count ≥ 10). Empirical data is compared against fitted power law (exponent α), log-normal, and truncated power law distributions. All three networks show power-law exponents in the typical biological range (α = 2.17–2.98), though log-normal and truncated power-law distributions provide statistically better fits. **(d–f)** Log-binned probability density of in-degree for each tissue confirming heavy-tailed hub structure.
